# Spectral imprint of structural embedding in effective connectivity

**DOI:** 10.1101/2025.06.19.660638

**Authors:** Matthew D. Greaves, Leonardo Novelli, James C. Pang, Alex Fornito, Adeel Razi

## Abstract

Neural fluctuations exhibit rich spectral profiles that reflects both local dynamics and structural (or anatomical) embedding. Yet, standard models of resting-state effective connectivity neglect structural embedding and assume uniformity in the timescales of regions’ endogenous fluctuations. We introduce a chromatic dynamic causal model (DCM) in which structural valency (or degree) modulates the spectral ‘color’ of endogenous fluctuations. Specifically, we assume a linear mapping between regional structural valency and the spectral exponent of scale-free auto-spectra. Simulations show this mapping can emerge as a generic consequence of structural embedding under minimal coupling in a non-equilibrium regime. We show chromatic DCM reliably recovers ground-truth parameters across network sizes and noise conditions, outperforming standard spectral DCM. Applied to empirical data, chromatic DCM reveals that valency–exponent mappings vary across a cortical hierarchy, and that its parameters are conserved across a homologous network in humans, macaques, marmosets, and mice. These findings advance a generative account of structure–function coupling and expand the repertoire of biophysical mechanisms available for inference in effective connectivity modeling.

## Main

To shed light on the mechanisms underlying observed neural dynamics, the neuroscience community has increasingly turned to both multimodal imaging and the modeling of effective connectivity^1,2^. For example, recently, the combination of structural (anatomical) and functional imaging has facilitated the discovery of a bipartition of single neurons’ receptive fields^3^, while altered patterns of directed influences between neuronal populations (effective connectivity) inferred from resting-state functional MRI (fMRI) have been shown to express early-stage dementia risk^4^. This work has largely progressed along parallel lines, however, and despite increasing interest in incorporating structural information into effective connectivity models^5^, such multimodal integration remains limited.

In this context, dynamic causal modelling represents a dominant framework for inferring resting-state effective connectivity, with its defining feature being the use of inversion of a generative (state-space) model to draw conclusions about unobserved variables that are thought to cause observed data^6,7^. Among model variants, the spectral dynamic causal model (DCM) is distinguished by its extensive validations^8^, which include in silico assessments of face and construct validity^9^, comparisons with invasive testing (using optogenetics^10^), multi-site reliability assessments^11^, and prediction of clinical outcomes^4,12^. From a practical standpoint, inversion in the spectral — rather than temporal — domain, where the model generates the expected cross-spectral density (CSD) of observed blood-oxygen-level-dependent (BOLD) responses, renders the inversion of large-scale models comprising dozens of regions tractable.

Its strengths notwithstanding, spectral DCM imposes a questionable simplification on the generative process: it assumes that fluctuations endogenous to neuronal populations — and specifically, their spectral profiles or ‘color’ — are identical for all regions^8^. This assumption is difficult to justify given that endogenous fluctuations likely reflect both local recurrent activity and interactions with neural elements beyond the modeled network, and that cytoarchitecture, connection density, and neuronal timescales exhibit robust and well-characterized regional variation^13,14^.

Indeed, empirical evidence motivates an alternative assumption. Research across species and imaging modalities indicates that a region’s structural valency (or degree) covaries with spectral properties of its intrinsic activity — namely, low-frequency power or the slope of its auto-spectrum (or power spectral density)^15–18^. These findings suggest that greater structural integration gives rise to ‘redder’ auto-spectra — consistent with slower, more autocorrelated dynamics — and support the view that this relationship arises, upstream of task-relevant neuronal computations, as a generic consequence of embedding within a non-equilibrium regime under minimal (global) coupling conditions^19^.

Here, we address these issues in a ‘chromatic’ DCM — so named in reference to the convention of labeling power spectra by color — in which structural valency shapes the spectral profile of endogenous fluctuations. Given empirical estimates of structural connectivity (Fig. 1a) and computed regional valency (Fig. 1b), the model assumes a monotonic relationship between normalized valency and the spectral exponent of a power-law model for auto-spectra (Fig. 1c). In the temporal domain, this translates to region-specific intrinsic timescales (Fig. 1d), upstream of both ensemble-level interactions (Fig. 1e) and the induction of hemodynamic responses (Fig. 1f).

**Fig. 1.**
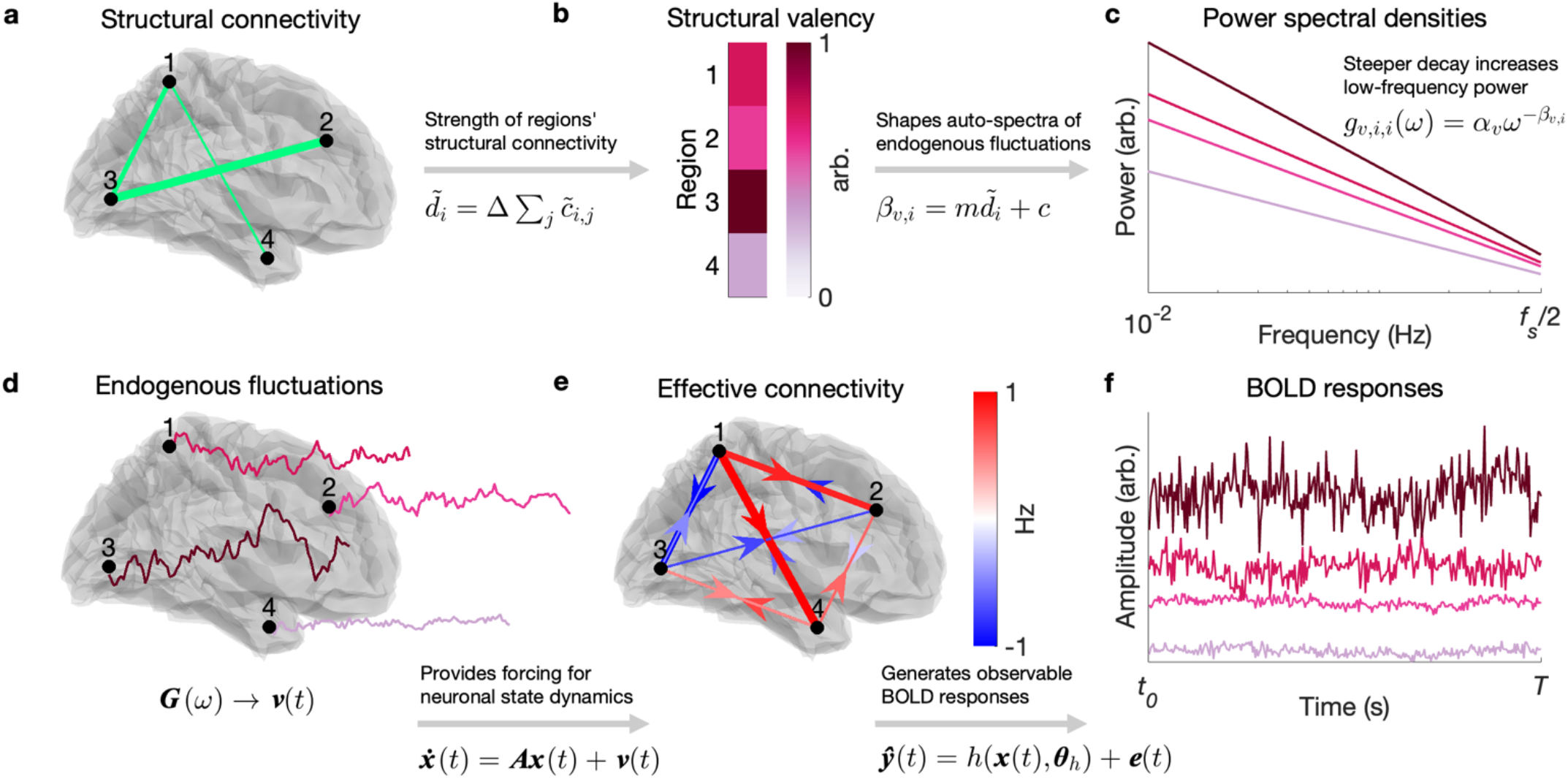
Chromatic DCM links structural embedding to spectral profiles of endogenous fluctuations. Panels illustrate the chromatic DCM framework, which incorporates regional anatomical embedding to constrain the spectral profile of endogenous fluctuations in the context of resting-state fMRI. White–burgundy color gradient reflects spectral color — from white (flat spectra) to pink and red (steeper, low-frequency dominated spectra) — and are used consistently across panels to denote region-specific properties. (**a**) Structural connectivity (with edge weight reflecting connection strength) is used to compute the normalized valency (degree) of each region, based on the total strength of its anatomical connections. (**b**) These valency values shape the spectral exponent of endogenous neuronal fluctuations via a linear mapping. (**c**) This results in region-specific auto-spectra (power spectral densities) for endogenous fluctuations, with a greater exponent corresponding to redder (steeper) spectra. (**d**) In the temporal domain, this corresponds to slower fluctuations in more structurally embedded regions. (**e**) These fluctuations drive neuronal dynamics upstream of effective connectivity. (**f**) Simulated BOLD responses show that (observed) timescales reflect both endogenous fluctuations and effective connectivity.

By formalizing these relationships, we address whether previously reported statistical regularities reflect deeper biophysical constraints. Using Monte Carlo simulations, we establish the feasibility of inferring this mechanism from data. We then show that a valency–spectra mapping arises in a minimalistic system of coupled oscillators operating near criticality, that it varies across a cortical hierarchy, and that its weighting of structural valency is conserved across homologous networks in humans, macaques, marmosets, and mice. Finally, we benchmark chromatic DCM against standard spectral DCM using simulated ground-truth models, showing that incorporating structural information improves the biophysical plausibility of resting-state effective connectivity estimates without sacrificing computational efficiency.

## Results

### A chromatic dynamic causal model

The chromatic DCM posits that structural valency shapes the timescales of endogenous neuronal fluctuations via a power-law spectral filter. The temporal-domain formulation assumes the following continuous-time state-space representation:

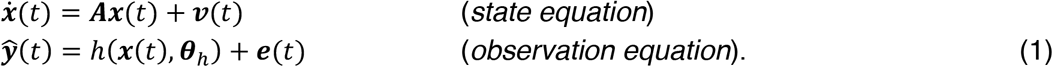

Here, in a system comprising *n r* neuronal populations, denotes the neuronal state vector, where each scalar function *x*_*i*_(*t*) ∈ ℝ describes the ensemble (or mean-field) activity of the *i*-th population at time *t*. The transition matrix 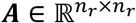 encodes the intra-and inter-regional modulation of the rates of change in this ensemble activity (quantifying the effective connectivity), in its diagonal and off-diagonal elements, respectively (Fig. 1e). In the observation equation, *h* is the hemodynamic response function (HRF) with free parameters ***θ***_*h*_ (Methods, Eqs. 4–5), which maps ensemble neuronal activity to expected BOLD responses ŷ (*t*) (Fig. 1f), which are compared to empirical data with parameterized residual precision (Methods, Eqs. 11–12). Per spectral DCM, both endogenous fluctuations ***x***(*t*), and observation error ***e***(*t*), are parameterized as power-law noise (Methods, Eq. 6); however, in the context of the model introduced here, the former is modulated by the brain’s structural architecture following:

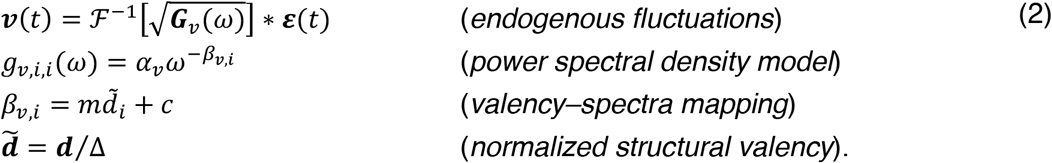

Here, *ℱ*^−1^ is the inverse Fourier transform, and 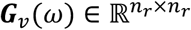 denotes a diagonal matrix-valued function of angular frequency *ω*, whose entries *g*_*v*,*i*,*i*_(*ω*) encode an auto-spectrum or power spectral density (PSD) model of region-specific endogenous fluctuations. The symbol ∗ denotes convolution, and *ϵ*(*t*)∼*N*(0, ***I***) is IID Gaussian noise. Under this formulation, the inverse Fourier transform of 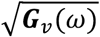 yields a temporal kernel that shapes the autocorrelation structure of ***v***(*t*). The PSD model is parameterized by a global amplitude *α*_*v*_ and a region-specific spectral exponent *β*_*v*,*i*_, which jointly determine the ‘color’ of the fluctuations: larger values of *β*_*v*,*i*_, yield redder (more temporally correlated) noise, while lower values yield whiter (less correlated) noise (Fig. 1c–d). Finally, spectral exponent *β*_*v*,*i*_ is linearly mapped from the normalized structural valency (or degree) 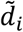, where *d*_*i*_ reflects the number or weight of structural connections associated with region *i*, and (per graph-theoretic notation) Δ is the maximal structural valency, *max*(***d***) (Fig. 1a–b). Note that in this valency–spectra mapping, the slope *m* and intercept *c* are constrained by two latent parameters such that *β*_*v*,*i*_ remains within a biologically plausible range (Methods, Eq. 3).

We highlight, here, that this formulation remains agnostic as to the choice of structural connectivity data: the normalized structural valency may — assuming non-negative values — be derived from weighted or binary (symmetric or asymmetric) matrices. Although the model is compatible with a range of data types (as demonstrated in later sections), our simulations predominantly focus on weighted symmetric structural connectivity, reflecting the current state of tractography methods, which typically assign weights to (undirected) streamline counts or connection probabilities.

### Network-dependent scaling of endogenous fluctuations

To establish data-informed priors for the latent parameters governing *m* and *c* in the valency–spectra mapping (λ_*c*_ and λ_*m*_, respectively; Methods, Eq. 3), we inverted subject-specific chromatic DCMs for 100 healthy adults from the Human Connectome Project (HCP^20^) using non-informative, low-precision priors. These first-level analyses utilized both resting-state fMRI data and structural connectivity derived from diffusion-weighted MRI and considered 20 effective connectivity networks defined by a hybrid atlas of 200 cortical regions (17 networks) and 32 subcortical regions (3 networks). In second-level analyses, posteriors for network-specific parameters were evaluated and aggregated using standard Bayesian modeling procedures implemented in the statistical parametric mapping (SPM) toolbox (Methods, Eqs. 13–16)^21^.

Consistent with prior work^22^, the baseline autocorrelation of endogenous fluctuations inferred for subcortical networks (as encoded by the intercept of the valency–spectra mapping) were lower than those for cortical networks (dashed box, Extended Data Fig. 1a). Furthermore, the extent to which structural valency modulated the autocorrelation of endogenous fluctuations differed between unimodal and transmodal cortical networks (Extended Data Fig. 1a), suggesting a hierarchical structure-function relationship^23,24^. In all subsequent analyses, the mean of across these aggregated network-specific posteriors furnished the prior expectations for valency–spectra mapping parameters (Extended Data Fig. 1b), with prior precisions determined by simulating 20 distinct hierarchical models (using network-specific posteriors shown in Extended Data Fig. 1a as the ground truth) and locating the precision settings that minimized the (average) root-mean-squared error (RMSE) between ground-truth and recovered parameters (Methods, Extended Data Fig. 1c).

### Recovery of ground-truth effective connectivity and valency–spectra mapping in silico

To establish the face validity of the chromatic DCM, we assessed the extent to which inverting the model successfully recovered parameters when the ground truth was known (simulated). Simulations were generated in the following manner. First, the normalized structural valency was derived from a random symmetric structural connectivity matrix (Fig. 2a) and mapped to spectral exponents using plausible valency– spectra mapping parameters (Fig. 2b, Supplementary Table 1). Then, region-specific, power-law endogenous fluctuations were modeled in the frequency domain (Fig. 1c) and projected into the time domain per Eq. 2 (Fig. 2d). Effective connectivity, modeled as random deviations from the simulated structural connectivity (Fig. 2e, Methods), determined neuronal state transitions (Fig. 2f), and these neuronal state variables yielded simulated BOLD responses via the HRF (Fig. 2g, Methods, Eqs. 4–5). To model realistic measurement conditions and assess robustness to noise, power-law observation error was added to each regional BOLD response at —unless otherwise noted — a signal-to-noise ratio (SNR) of 0 dB (equal signal and noise power; Fig. 2h, Methods, Eq. 19).

**Fig. 2.**
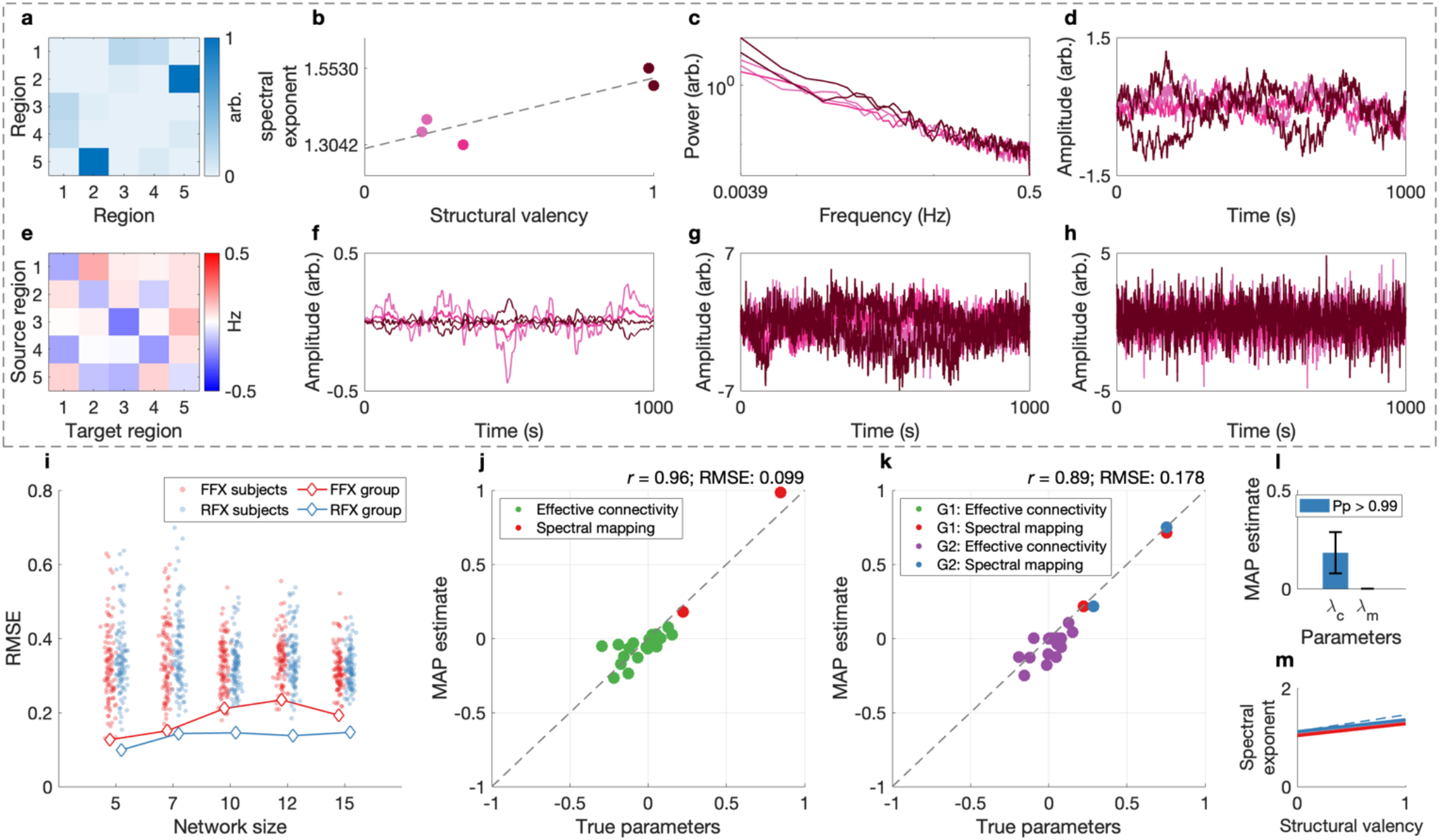
Accurate recovery of effective connectivity and valency–spectra mapping in silico. Panels within dashed box illustrate the process used to simulate data for a five-node network (white-through-burgundy colors consistent with region identity and spectral color); panels outside dashed box show results after inverting the chromatic DCM on the simulated data. (**a–d**) Simulated endogenous fluctuations were shaped by structural valency (**a–b**), with region-specific auto-spectra (**c**) and time-domain signals (**d**) utilized in simulations reflecting these auto-spectral profiles. (**e–g**) Random effective connectivity (**e**) and endogenous fluctuations drove neuronal dynamics (**f**) and synthetic BOLD responses (**g**), with observation noise added (**h**). (**i**) Across five network sizes and two modeling regimes (FFX, red; RFX, blue), the accuracy of parameter recovery remained consistent as indicated by RMSE (with RFX outperforming to FFX modeling). RMSE is shown for both subject-and group-level models (diamonds do not represent average RMSE across subjects). (**j**) True and estimated parameters (RFX, 5-node model) showed high agreement for both effective connectivity (green) and valency–spectra mapping (red). Parameters are log-transformed to aid visualization. (**k**) Group differences in valency–spectra mapping was detected in a second RFX simulation, with no spurious differences in effective connectivity. Parameters are log-transformed to aid visualization. (**l**) Posterior probability of valency–spectra mapping differences exceeded 0.99. (**m**) Group-level MAP estimates reproduced the correct direction of the small valency–spectra difference, although slope-related differences were attenuated (dashed lines show ground truth).

To examine the impact of network size on parameter recovery, we simulated BOLD responses for 100 synthetic subjects using network models of varying size (*n*_*r*_ = {5,7,10,12,15}) and two distinct second-level modeling regimes. In the fixed-effects (FFX) simulations, all subjects shared identical ground-truth parameters, with variability arising solely from stochastic endogenous fluctuations (and, consequently, BOLD responses and observation error; Methods). In the random-effects (RFX) simulations, subject-specific parameters were sampled from a group-level distribution centered on the same ground-truth values used in the FFX simulations, thereby introducing inter-subject variability in both effective connectivity and valency– spectra mapping (Methods, Supplementary Table 1). Chromatic DCMs were inverted separately for each network size and modeling approach, with group-level inference conducted using Bayesian model averaging (BMA) in the FFX case and parametric empirical Bayes (PEB) in the RFX case (Methods, Eqs. 13–16)^25,26^.

As shown in Fig. 2i, RFX inference consistently outperformed FFX inference across all network sizes, as indicated by lower RMSE values when comparing the true and maximum a posteriori (MAP) effective connectivity and valency–spectra mapping parameters (blue versus red scatters and line plot). Notably, across both modeling regimes, estimation accuracy remained stable as a function of network size, suggesting that the chromatic DCM supports robust parameter recovery in high-dimensional settings. Fig. 2j shows a parity plot for the five-region RFX group model, comparing true and MAP parameter estimates across both effective connectivity (green) and valency–spectra mapping parameters (purple). Parameter estimates closely follow the identity line (dashed), with a high correlation and low RMSE.

Next, to assess the separability of model components, we simulated BOLD responses under a five-region network model for a second group of 100 synthetic subjects. In this RFX simulation, subjects shared the same underlying distribution of effective connectivity as the first group (Fig. 2j), but valency–spectra mapping parameters were drawn from a distinct group-level distribution (parameters per Supplementary Table 1, with the exception that *m* = 1/4). This design allowed us to identify whether group differences in valency–spectra mapping could be correctly inferred without spurious differences emerging in effective connectivity estimates. Inversion of chromatic DCMs and estimation of a second-level PEB model comprising DCMs from both groups revealed group differences in the parameters governing the valency–spectra mapping, evident in the distance between MAP estimates in Fig. 2k (Methods, Eqs. 15–16). This difference was significant, with the posterior probability that the relevant group difference parameters were non-zero exceeding 0.99 (Fig. 2l), while no significant differences were observed in effective connectivity. Finally, although the MAP estimates captured the correct direction of the effect (Fig. 2m), the difference was attenuated relative to ground truth.

### The valency–spectra mapping emerges in a metastable regime

To investigate whether a minimalistic coupled system induces a mapping from structural valency to spectral exponents per Eq. 2, we simulated dynamics over both random and empirical structural connectivity using a network of coupled Hopf oscillators (a canonical dynamical system in neuroscience, utilized in modeling both the transition to oscillatory activity in single neurons, and characterizing whole-brain resting-state dynamics)^27,28^. Note that here, we binarized structural connectivity to model unitary coupling relationships between neural elements, independent of spatial scale. In addition, we utilized a single global (or background) gain term, ensuring that observed spectral variation reflected intrinsic embedding in the structural network, rather than node-specific tuning.

The behavior of the Hopf bifurcation model is governed by a bifurcation parameter *μ* (Methods, Eq. 20), which determines whether each oscillator tends toward a fixed point (damped dynamics) or a limit cycle (sustained oscillations). Fig. 3a illustrates this behavior in a two-dimensional phase space, with the horizontal and vertical axes corresponding to the real and imaginary components of the oscillator’s state, respectively. Arrows show the direction and rate of change at discrete points in the state space while the spiral trajectory illustrates how the system evolves over time from a given initial condition. Here, with *μ* = −0.001, the system sits just below the bifurcation (μ = 0), producing slowly decaying oscillations (a near-critical regime characterized by increased susceptibility to inputs).

**Fig. 3.**
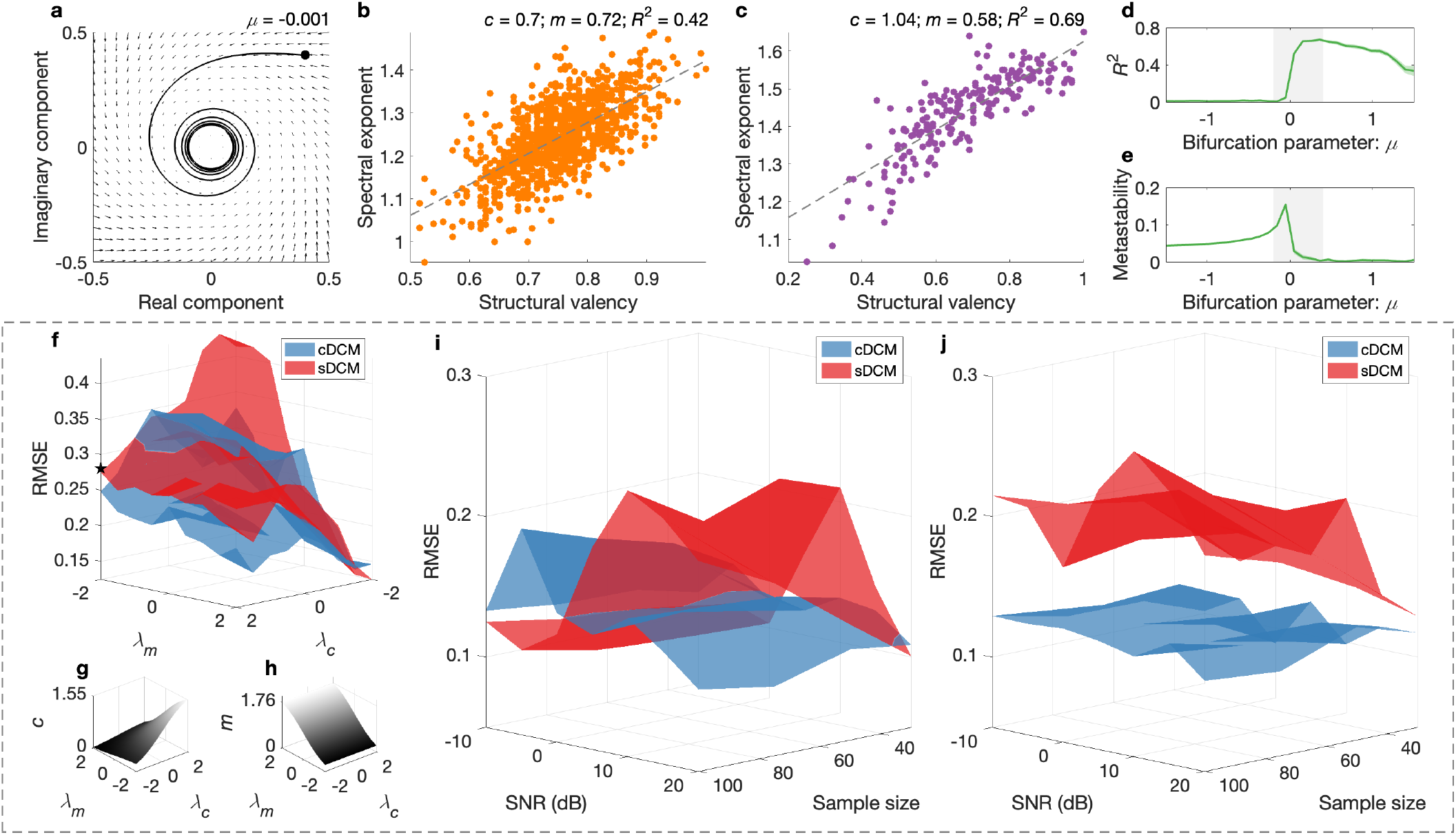
Valency–spectra mapping emerges in a metastable regime and supports inference in dynamic causal modeling. Panels above dashed box illustrate how a valency–spectra mapping arises in a near-critical dynamical regime; panels within dashed box benchmark chromatic DCM (cDCM) against spectral DCM (sDCM) in terms of parameter recovery and robustness. (**a**) A Hopf oscillator with bifurcation parameter *μ* = −0.001 shows damped oscillatory behavior. (**b–c**) In simulations using random (**b**) and empirical (**c**) structural connectivity, regional spectral exponents were linearly related to structural valency. (**d–e**) The goodness of fit (**d**) of this relationship peaked near the bifurcation point where metastability (**e**) was maximal (indicated by gray region), defined as variability in global phase synchrony (bootstrapped 95% confidence envelopes shown). (**f**) Across a grid of valency–spectra mapping parameters (*λ*_*c*_, *λ*_*m*_), chromatic DCM showed lower RMSE than spectral DCM (pentagon shows parameter settings used in later panels), particularly when endogenous fluctuations were white-like (**g–h**). (**i**) Chromatic DCM maintained lower RMSE across increasing SNR and sample size, whereas spectral DCM performance degraded. (**j**) When observation noise was made more autocorrelated (*β*_*e*_ = 1), the chromatic DCM advantage over spectral DCM increased.

In the context of this near-critical regime, we first examined whether a regions’ structural valency predicted the spectral exponent of its simulated activity. As shown in Fig. 3b, for random structural connectivity comprising *n*_*r*_ = 1000 nodes, a clear positive linear relationship emerged (coefficient of determination, *R*^2^ = 0.42), with higher-valency nodes exhibiting slower fluctuations. The same analysis using empirical structural connectivity (*n*_*r*_ = 200) revealed a stronger relationship (Fig. 3c, *R*^2^ = 0.69). Fig. 3d–e show that, in analyses where we varied the bifurcation parameter *μ* ∈ [−1.5, 1.5] and utilized random structural connectivity (*n*_*r*_ = 200), the correspondence between the valency and spectral exponent peaked in a near critical region (near the bifurcation point): *R*^2^ between valency and spectral exponent was highest (Fig. 3d) in the same region of the parameter space where metastability was maximized (Fig. 3e). Here, metastability was defined as the standard deviation of the Kuramoto order parameter, which indexes the degree of global phase synchrony over time (Methods, Eq. 21).

### Benchmarking construct validity against spectral dynamic causal modelling

Chromatic DCM represents an extension of the well-known spectral DCM. The main differences between spectral DCM and the chromatic DCM presented here is that the latter is multimodal, and that spectral DCM — per its canonical implementation — assumes a single PSD model of endogenous fluctuations for all network regions (and, in the limit, all brain regions), with *g*_*v*,*i*,*i*_(*ω*) = *g*_*v*,*j*,*j*_(*ω*) (Eq. 2)^8^. Thus, spectral DCM serves as the most natural comparator model for assessing the construct validity of chromatic DCM.

In Fig. 3 (dashed box), we focus on the recovery of effective connectivity in the context of RFX simulations. First, we simulated BOLD responses for 100 synthetic subjects under a five-region network across a grid of plausible valency–spectra mapping parameters, with *λ*_*c*_, *λ*_*m*_ ∈ {−2, −1.5, . . ., 2} and all other parameters set per Supplementary Table 1. Chromatic and spectral DCMs were then inverted and summarized at the group level using PEB, and the true and MAP estimates of effective connectivity were compared across the parameter grid. Fig. 3f shows that the RMSE was generally lower for chromatic DCM, and that the accuracy of spectral DCM was poorest in regions of the parameter space where both the intercept and slope of the valency–spectra mapping were near zero, and endogenous fluctuations were noisiest or most ‘white’ (transformed parameters, Fig. 3g–h).

To assess the robustness of group-level inference across chromatic and spectral DCMs when both data quality and sample sizes are variable, we simulated BOLD responses for 100 synthetic subjects under a five-region network across a grid of SNRs (SNR ∈ {−20, −15, . . ., 20}) and sample sizes (*S* ∈ {30,50,75, 100}). In these simulations, *λ*_*c*_ = 2 and *λ*_*m*_ = −2 (black pentagram, Fig. 3f), with all other parameters set per Supplementary Table 1. These parameter settings correspond to a scenario in which endogenous fluctuations are uniformly autocorrelated — the valency–spectra mapping has a negligible slope (Fig. 3g–h) — making it a fair test case where spectral DCM is expected to perform well. Fig. 3i shows that, while RMSE for chromatic DCM generally improved with increasing SNRs and sample size, the RMSE for spectral DCM paradoxically worsened under the same conditions.

To investigate whether this difference in accuracy was influenced by the autocorrelation structure of the observation error, we repeated the simulation and analysis shown in Fig. 3i, altering the global spectral exponent for the observation error PSD model (Methods, Eq. 6), from *β*_*e*_ = 1/3 to *β*_*e*_ = 1, consistent with noise arising from self-similar physiological processes (for example, respiration or head motion), and equivalent to the prior expectation for *β*_*e*_ in both chromatic and spectral DCMs. Fig. 3j shows that this alteration exacerbated the difference in accuracy between models: RMSE for spectral DCM worsened across the SNR and sample size grid, whereas RMSE for chromatic DCM remained stable.

### Cross-species validity of the chromatic dynamic causal model

To assess the generalizability of chromatic DCM estimates across species, we applied the model to a homologous brain circuit in humans (n = 100), macaques (n = 5), marmosets (n = 13), and mice (n = 193). For humans, structural connectivity was derived from tractography applied to diffusion-weighted MRI, and resting-state fMRI was acquired during normal waking conditions^20^. For non-human animals, ground-truth structural connectivity was obtained from invasive tract-tracing data^29–31^, and, with the exception of marmosets (Methods)^32^, resting-state fMRI data were recorded under anesthesia^33,33–35^. In the macaque dataset, the experimenter-defined parcellation yielded data for 29 macroscopic brain regions, averaged across hemispheres. This allowed us to invert whole-brain chromatic DCMs (within feasible computational limits) under three distinct definitions of structural valency — based on efferent (outgoing), afferent (incoming), or average connection strengths. We then compared these model variants in terms of their free energy and evaluated the corresponding estimates of effective connectivity by comparing them to the asymmetric whole-brain structural connectivity matrix.

Four homologous regions were selected based on established anatomical correspondences across species and the availability (in the case of non-human animals) of regional tract-tracing data: primary visual cortex (V1), middle temporal area (V5), primary motor cortex (M1), and anterior cingulate cortex (ACC; Fig. 4a–d, Supplementary Material). For each species, subject-specific chromatic DCMs were inverted using regional BOLD time series data and connection-averaged structural valency, with effective connectivity and the valency–spectra mapping summarized at the group level using PEB (Fig. 4e–h). As shown in Fig. 4i, group-level MAP estimates of effective connectivity were highly consistent across species, as assessed via pairwise correlation. While baseline spectral exponents differed across species — implying variation in the intrinsic autocorrelation of fluctuations in less structurally embedded regions — the degree to which structural valency modulated endogenous fluctuations was well aligned, as reflected in the slope of each valency–spectra mapping (Fig. 4j).

**Fig. 4.**
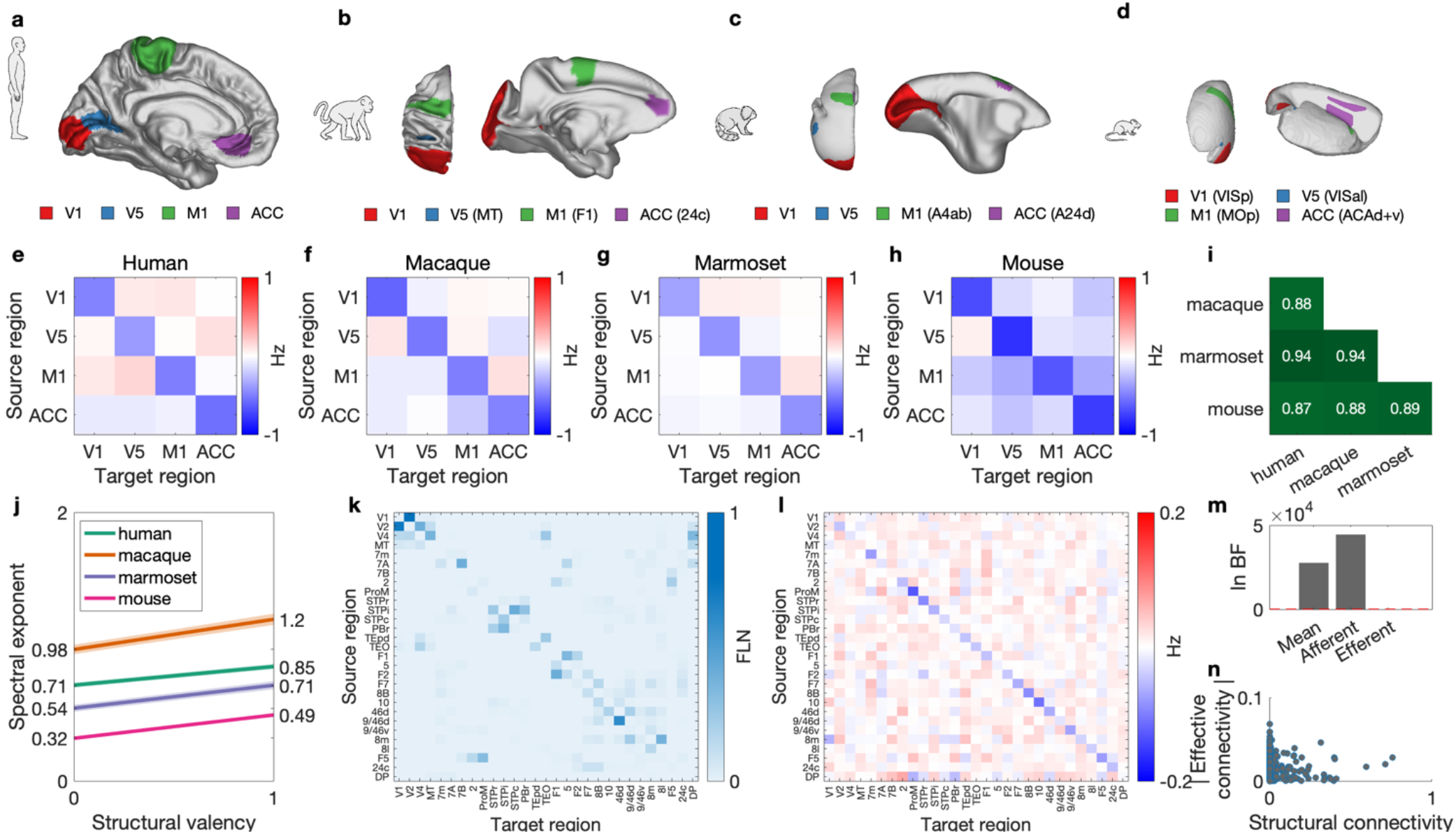
Cross-species validity of chromatic DCM in homologous brain circuits. Panels (**a–d**) show homologous cortical regions (V1, V5, M1, ACC) across human (**a**), macaque (**b**), marmoset (**c**), and mouse (**d**) brains, used to evaluate chromatic DCM estimates across species. Panels show relevant regions on sagittal views of left hemisphere surfaces, with posterior views included where they provide additional context. Species-specific (atlas) nomenclature is indicated in parentheses (Supplementary Table 3). (**e–h**) Group-level effective connectivity matrices, estimated using chromatic DCM and MRI-or tract-tracing based structural connectivity, reveal conserved profiles across species. (**i**) Pairwise correlations of effective connectivity matrices indicate strong cross-species similarity (all > 0.87). (**j**) Valency–spectra mappings reveal consistent weighting of endogenous fluctuations by structural valency. (**k**) Macaque whole-brain structural connectivity matrix derived from invasive tract-tracing (FLN: fraction of labeled neurons), averaged across hemispheres and symmetrized. (**l**) Corresponding effective connectivity matrix estimated using chromatic DCM with afferent valency. Inhibitory self-connections dominate the diagonal (log-scaled for visualization). (**m**) Comparison of model evidence across macaque whole-brain models, each defined using a different computation of valency (Mean: symmetrized connectivity). Bars show the log Bayes factor relative to the lowest-evidence model. Red dashed line marks a threshold of 3, corresponding to 20× greater evidence for the comparator model. (**n**) Relationship between structural and absolute effective connectivity in the macaque dataset, showing that directed effective coupling does not trivially mirror structural coupling strength.

Additional results from the macaque whole-brain analysis are presented in Fig. 4k–n. The tract-tracing-derived structural connectivity matrix (Fig. 4k) shows characteristic asymmetries, which provided the basis for the computation of efferent, afferent, and mean (symmetrized) structural valency used in model inversion. The corresponding group-level whole-brain effective connectivity estimates obtained using afferent valency (Fig. 4l), provided substantially higher model evidence than alternatives (Fig. 4m), with a log Bayes factor well beyond the conventional threshold (red dashed line, Fig. 4m). Finally, Fig. 4n shows that absolute effective connections do not trivially mirror the fraction of labeled neurons (FLN, Fig. 4k).

## Discussion

Empirical evidence indicates that a region’s structural valency covaries with spectral properties of its intrinsic activity, challenging a key assumption of the most well-known model of resting-state effective connectivity.

Our chromatic DCM accounts for this relationship by leveraging multimodal imaging and formalizing a mechanism whereby valency shapes the color — or autocorrelational structure — of the auto-spectra of endogenous fluctuations. We establish the feasibility of inferring this mechanism from BOLD responses and identify continuities with prior work: its parameterization differs across both cortical–subcortical distinctions and an approximate unimodal–transmodal heirachy, and — within a network of homologous regions — the slope of the mapping is conserved across mammalian species. Our view of this mechanism as arising as a general consequence of minimal coupling was supported by analyses in which applications of the Hopf model exhibited similar structure–spectra relationships. Additionally, in analyses of construct validity, by simulating scenarios where spectral DCM is expected to perform well, we show that chromatic DCM yields more plausible estimates of effective connectivity. The chromatic DCM can be applied to fMRI data and variously defined structural connectivity data and opens new possibilities for examining both structure–function relationships and effective connectivity in future work.

Although previous studies have incorporated structural information into models of directed connectivity, our work demonstrates the feasibility of inferring a structure–function mechanism encoded upstream of macroscopic interregional communication. Most existing approaches to integrating structural connectivity into directed connectivity treat binarized structural connectivity as a mask or skeleton upon which putative directed influences play out^36,37^. Others integrate structural information by specifying priors over state-transition weights^38,39^, typically under the assumption that stronger structural connectivity implies a broader (higher variance) zero-mean prior for the corresponding directed connection. While this regularization-based approach has proved useful — increasing model evidence and yielding reliable structure-to-prior-variance relationships — it remains heuristic^5^. That is, in contrast to the modeling assumption, greater white matter integrity between two regions should, if anything, imply more precise (not more variable) interactions^40^. That said, our findings motivate future work integrating this pragmatic assumption into the inference of effective connectivity in the context of chromatic DCM.

By relating structural connectivity to the auto-spectra of endogenous fluctuations, our model expands the repertoire of causal mechanisms over which one can conduct inference within the framework of dynamic causal modelling. By simulating this relationship using a dynamical system in which the metastability of dynamics is parametrically controlled, we lend support to the idea that the valency–spectra relationships observed here emerge from a region’s embedding within a system operating near the edge of a bifurcation. Establishing the parameter ranges, input conditions, and dynamical constraints under which this behavior emerges — and determining whether it can be recapitulated using an autoregressive linearization of the same system — will be a focus of future work.

In the context of chromatic DCM’s application to a homologous network across mammalian species, we find that both the effective connectivity and the valency–spectra relationship exhibit consistency. In the latter case, although the baseline auto-spectral properties of endogenous fluctuations differ across species, the extent to which structural valency modulates these fluctuations is well aligned. This suggests that the mapping may conserved within homologous networks across evolutionarily distant systems. Whole-brain analyses in macaques suggest that modeling spectral exponents as a function of afferent (incoming) connectivity yields superior model evidence, supporting the view that endogenous fluctuations reflect the passive (or task-free) computation of external influences, rather than locally generated activity alone.

Taken together, by leveraging multimodal imaging and the modelling of effective connectivity, our findings highlight a biophysically plausible structure–function mapping that may underlie the spectral architecture of resting-state neural dynamics. By embedding this mapping within a chromatic DCM, we expand the class of effective mechanisms over which one can conduct inference in the context of dynamic causal modelling. Noting that we employed PEB under the a priori assumption that the likelihood’s precision was identical to that assumed under chromatic DCM, future work will be necessary to refine model inversion procedures for truthful characterization of group-level differences. Further directions include exploring the behavioral and clinical relevance of the inferred valency–spectra mappings and their downstream influence on structurally informed effective connectivity. This will clarify whether these parameters provide useful markers of health and disease.

## Methods

### Completing the chromatic dynamic causal model

To complete the specification of the chromatic DCM (Eqs. 1–2), we detail the latent constraints on the valency–spectra mapping, the form of the HRF, the form of the observation error, and the model’s projection into the spectral (frequency) domain. First, to ensure that the valency–spectra mapping (Eq. 2) did not yield PSD models that were biologically implausible or implied nonstationary processes^22,41^, its slope *m* and intercept *c* were constrained following:

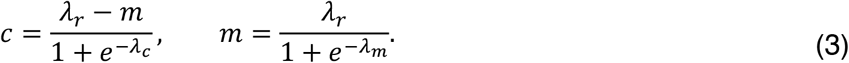

Here, *λ*_*r*_ is the maximum possible value of *β*_*i*_, which was fixed at 2 in all analyses. The parameters *λ*_*c*_ and *λ*_*m*_ were treated as latent variables with Gaussian priors, whose means and variances were determined empirically (Extended Data Fig. 1b, Supplementary Table 2). Except for analyses aimed at establishing these data-informed priors, in all analyses, after inverting DCMs (and prior to any group-level inference), posterior uncertainty in *λ*_*c*_ and *λ*_*m*_ was propagated to the transformed parameters *c* and *m* using the delta method.

In Eq. 1, *h*(*x*(*t*), ***θ***_*h*_) represents the well-known ‘Balloon’ HRF that yields expected BOLD responses from regional neuronal states *x*_*i*_(*t*) as follows^42^:

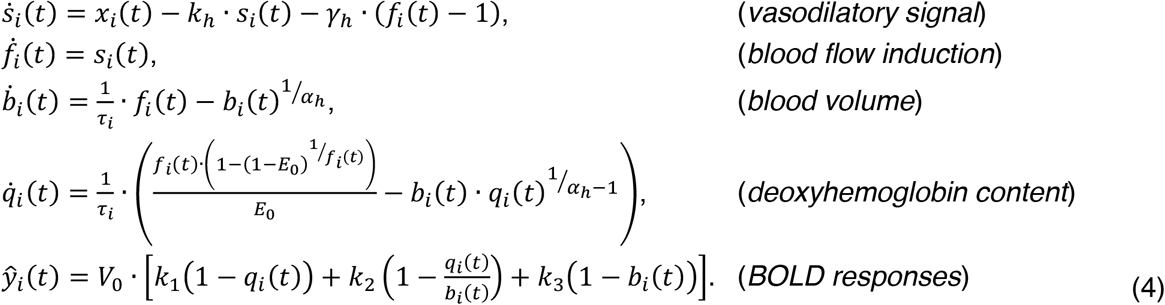

Here, neuronal activity *x*_*i*_(*t*) drives a vasodilatory signal *s*_*i*_(*t*), which increases blood flow *f*_*i*_(*t*), expands blood volume *b*_*i*_(*t*) via Grubb’s law, alters deoxyhemoglobin content *q*_*i*_(*t*) through oxygen extraction, and together these variables determine the expected BOLD response ŷ_*i*_(*t*). The vasodilatory signal decays at rate *k*_*h*_, the feedback of blood flow regulated by ***γ***_*h*_ = 0.32, ***τ*** = [*τ*_1_,…, *τ*_n_] are region-specific transit times, *α*_*h*_ = 0.32 is Grubb’s (vessel stiffness) exponent, and *E*_0_ = 0.4 and *V*_0_ = 4 represent the resting oxygen extraction fraction and resting venous blood volume fraction, respectively. In the last line of Eq. 4, coefficients *k*_1_, *k*_2_, and *k*_3_ represent the contributions of the intra-and extra-vascular compartments to the BOLD response:

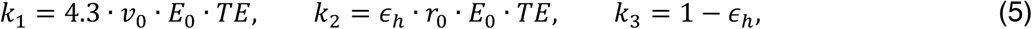

where *v*_0_ = 40.3 is the frequency offset at the outer surface of magnetized vessels, *TE* = 0.04 is the echo time, *r*_0_ = 25 is the slope of the intravascular relaxation rate, and *ϵ*_*h*_ is the ratio of intra-to extra-vascular signal contributions. The set of parameters that were free to vary is given by ***θ***_*h*_ = {*k*_*h*_, *τ, ϵ*_*h*_}.

Regarding the model of the observation error ***e***(*t*) (Eq. 1), like the PSD model for region-specific endogenous fluctuations (Eq. 2), observation error for the *i*-th neuronal population was assumed be power-law:

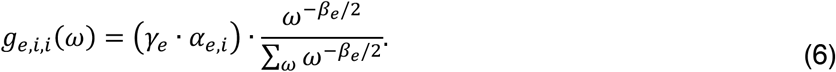

Here, 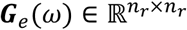 is a diagonal matrix, and the PSD model was controlled by a global spectral exponent *β*_*e*_, and region-specific amplitude parameter *α*_*e*,*i*_ normalized by a global factor ***γ***_*e*_.

Finally, using the Fourier transform, *ℱ*, we projected chromatic DCM into the spectral domain, such that it generated the expected cross-spectral density (CSD) of BOLD responses Ĝ *y*(*ω*) = [*ℱ*{ŷ +(*t*)}*ℱ*{ŷ +(*t*)}^*†*^], where † denotes the conjugate transpose. Putting this all together, the spectral equivalent of Eq. 1 reads:

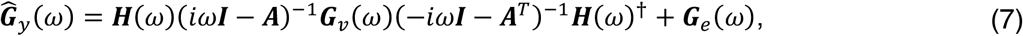

where ***H***(*ω*) is the Fourier transform of the HRF (Eqs. 4–5). Here, notably, the latent neuronal state in the frequency domain, ***X***(*ω*), has been factored out via the substitution ***X***(*ω*)***X***(*ω*)^*†*^ = (*iω****I*** − ***A***)^−1^*G*_*v*_(*ω*)(−*iω****I*** − ***A***^*T*^)^−1^. For a didactic introduction to these derivations, see Novelli and colleagues^43^.

To construct the empirical CSD matrix *G*_*y*_(*ω*) for model inversion, we fitted a multivariate autoregressive model of order *p* = 8 to observed BOLD time series using the variational Bayesian procedure described by Penny and Roberts^44^. This procedure yielded a set of autoregressive coefficient matrices 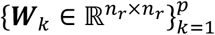 and a residual covariance matrix 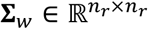. The standard spectral factorization then produced *G*_*y*_(*ω*) as:

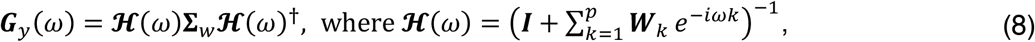

Consistent with prior related work, ***G***_*y*_(*ω*) was estimated at *n*_*f*_ = 32 linearly spaced frequencies between 1/128 Hz and the Nyquist frequency: 1/2. *TR*, where *TR* is the repetition time (sampling interval)^45^.

In the following sections, we refer to Eq. 7 as the first-level model 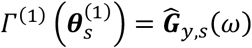, where *Γ*^(1)^ is a given DCM architecture, the subscript *s* = 1, . . ., *S* indexes subject-specific parameters and data, and the superscript (1) distinguishes these elements from higher-order quantities. Unless otherwise specified, we use 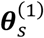 to refer to all subject-specific parameters.

### Inverting the chromatic dynamic causal model

With priors specified for chromatic DCM parameters (Supplementary Table 2), the model can be cast as a generative model (in the statistical sense). Namely, it provides a model of the joint probability distribution over observed data *G*_*y*,*s*_ and model parameters 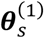, given by: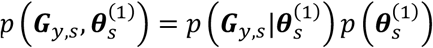, where 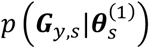 is the likelihood implied by the model, and 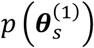 is the prior distribution over parameters^8^. Inverting the model — inferring 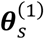 from observed data — thus corresponds to computing the posterior distribution 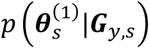, which follows directly from Bayes’ rule, expressed here in generic form (with data *y* and model parameters ***θ***):

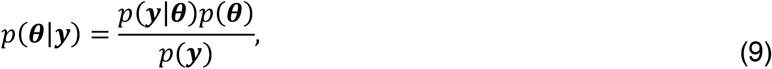

Given that evaluating the model evidence *p*(***y***) is generally intractable, here, we employed variational Bayes under the Laplace approximation (VBL) to approximate posterior distributions (as implemented in the SPM toolbox)^46^. This approach assumes that the true posterior distribution is Gaussian around its mode. The fundamental VBL equation is:

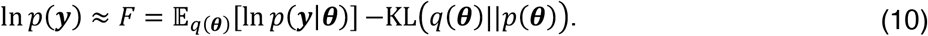

Here, the free energy *F* is a variational approximation of the log-model evidence, *E*_*q*(***θ***)_[ln *p*(*y*|***θ***)] is the expected log-likelihood under a variational — approximate posterior — distribution *q*(***θ***), and KL*vq*(***θ***)||*p*(***θ***)w is the Kullback–Leibler divergence between this approximate posterior distribution *q*(***θ***), and the prior *p*(***θ***). This formulation permits the optimization of *q*(***θ***) to maximize free energy— thereby balancing model accuracy and complexity — via a procedure described in detail by Zeidman and colleagues^47^.

In the context of chromatic DCM, the expected log-likelihood *E*_*q*(***θ***)_[ln *p*(*y*|***θ***)], quantifies the fit between the expected and observed CSDs under the approximate posterior. This is expressed as:

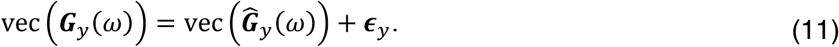

In the context of VBL, the residual error ϵ_*y*_∼*N*(**0, Σ**_*y*_) is parameterized in terms of its precision. Per the implementation utilized here, the residual error precision 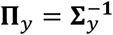, is expressed as a weighted sum of 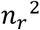 precision components:

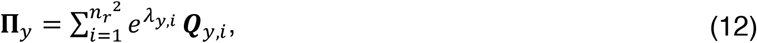

where 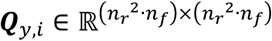 is a sparse, block-diagonal matrix with an *n*-dimensional identity sub-block targeting the *i*-th region pair (including self-connections). Here, the precision weights *λ*_*y*,*i*_ are drawn from a hyperprior, whose hyperparameters we determined in silico (Methods, Extended Data Fig. 1d-e).

### Group-level inference procedures

In FFX analyses (for example, Fig. 2i) and analyses of data-informed priors (Extended Data Fig. 1), we employed BMA to aggregate parameters across subjects and models, respectively^48^. In the case of the former analyses, under the assumption that all (simulated) subjects share a common model structure *Γ*^(1)^, and common parameterization, group-level parameters were obtained via simple parameter averaging across subjects’ posteriors:

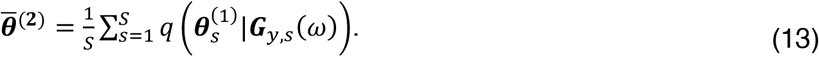

In the case of the latter analyses (Extended Data Fig. 1), aggregate group-level posterior distributions were computed by aggregating posterior estimates across a grid of ***K*** second-level models (denoted 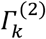), each differing in prior assumptions. These posteriors were weighted by their corresponding posterior model probabilities *p*_*k*_, which are computed via a softmax transformation of the free energy *F*_*k*_ associated with each model. Thus, we obtained an evidence-weighted aggregate of second-level parameters 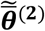, via:

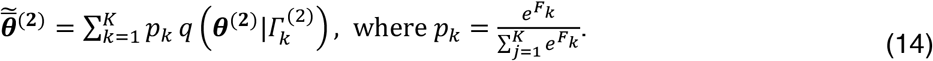

In contrast, for RFX analyses (for example, Fig. 2j–k), we used PEB to model between-subject variability via a second-level Bayesian general linear model^25^. Vertically stacking the vectorized MAP estimates of interest for *S* subjects, 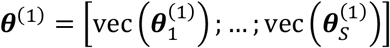, permits the following hierarchical formulation:

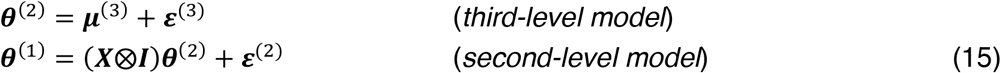

Here, at the third level, ***θ***^(2)^ ∈ ℝ^*D*^ are group-level parameters modeled as deviations ***ε***^(3)^∼*N*(0, **Σ**^(3)^) from prior expectations ***μ***^(3)^, and *D* is the cardinality of 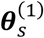. At the second level, the design matrix ***X*** ∈ ℝ^S*×k*^ (where *k* is the number of regressors, including any intercept) and Kronecker product ***X***⨂***I*** (where the identify matrix ***I*** is *D*-dimensional) ensure that ***θ***^(2)^ is appropriately tiled to each parameter in ***θ***^(1)^ ∈ ℝ^S*D*^. Following conventions, ***μ***^(3)^ and **Σ**^(3)^ duplicated the prior mean and covariance, respectively, of the prior distribution over 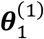, and the residuals *ϵ*^(3)^∼*N*(0, **Σ**^(3)^) were parametrized in terms of a scaled precision component:

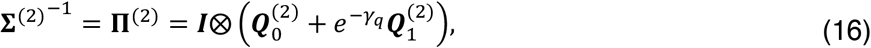

where ***I*** is an *S*-dimensional identity matrix, 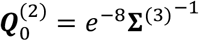 is the lower-bound precision, 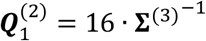 and the precision scale parameter ***γ***_*q*_ is inferred from the data (Supplementary Table 2).

With these elements specified, Eq. 14 can be naturally cast in terms of Gaussian densities and, in turn, as a generative model: *pv****θ***^(1)^, ***θ***^(2)^w = *pv****θ***^(1)^|***θ***^(2)^w*pv****θ***^(2)^w. Per first-level analyses, in all RFX analyses Eq. 14 was inverted using VBL (Eq. 9).

### Establishing data-informed priors for valency–spectra mapping parameters

To establish data-informed priors for *λ*_*c*_ and *λ*_*m*_, which govern the valency–spectra mapping (Eq. 2), we inverted subject-specific chromatic DCMs for 100 healthy adults from the HCP under non-informative, low-precision priors for *λ*_*c*_∼*N*(− ln 255, 1) and *λ*_*m*_∼*N*(− ln 255, 1), whose means corresponded to *c* = *m* = 1/128. With the additional exception of hyperparameter settings — set to the default for spectral DCM, *λ*_*y*,*i*_∼*N*(8, 1/128) — all priors were specified per Supplementary Table 2.

First-level model inversion utilized both resting-state fMRI data and structural connectivity derived from tractography applied to diffusion-weighted MRI and considered 20 effective connectivity networks. In second-level analyses, per network, first-level parameters 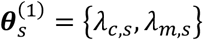, were taken to the second level, and PEB models were inverted, yielding group-level posteriors per network. Next, we explored candidate prior expectations and variances for *λ*_*c*_ and *λ*_*m*_ via grid search and Bayesian model reduction (BMR), an analytic means of computing the approximate posterior and free energy for alternative (reduced) models from an inverted full (parent) model (by adjusting the sufficient statistics of the prior distribution)^25^. Prior means were varied from −5 to 5 in integer steps, and prior precisions were varied as powers of 2, with integer exponents ranging from 2 to 7 (that is, from 2^2^ to 2^7^). This resulted in a 4-dimensional grid spanning all combinations of expectations and variances for the two latent parameters, and a total of 4,356 reduced models per network.

Per network, reduced models were aggregated using BMA, which provided an evidence-weighted aggregate valency–spectra mapping per network (Extended Data Fig. 1a). The mean of these network-specific aggregate exponent mapping parameters (rounded to the nearest tenth) then furnished the prior expectations for *λ*_*c*_ and *λ*_*m*_, utilized in all subsequent analyses (Extended Data Fig. 1).

To select appropriate prior precisions for *λ*_*c*_ and *λ*_*m*_, we simulated BOLD responses for 50 synthetic subjects under 20 distinct five-region network models that replicated both features of the HCP dataset and the results reported in Extended Data Fig. 1. Namely, in simulations, we used the network-specific evidence-weighted aggregate parameters for the valency–spectra mappings as ground truth values for *c* and *m* (Extended Data Fig. 1a), and reflecting the HCP’s high data quality, set *T* = 1200, *TR* = 0.72, and SNR = 20 (all other parameters per Supplementary Table 1). Then, rather than re-running simulations for various precision combinations, we inverted the 1000 chromatic DCMs under priors *p*(*λ*_*c*_)∼*N*(− ln 4/5, 1) and *p*(*λ*_*m*_)∼*N*(− ln 1/9, 1), inverted the 20 network-specific PEB models, and leveraged BMR to re-evaluate each model under different prior precisions (ranging, again, as powers of 2, with integer exponents ranging from 2 to 7).

For each prior precision setting, we computed the RMSE between the true and MAP estimates of *c* and *m*. The precisions that minimized the average RMSE across the 20 network models — results shown in Extended Data Fig. 1c — were selected for use in all subsequent analyses (standard error envelope, Extended Data Fig. b), to ensure that the priors were sufficiently flexible to capture diverse valency-to-spectral exponent relationships.

### Establishing residual error hyperparameters in silico

To identify hyperprior settings in chromatic DCM, we conducted a systematic grid search over combinations of prior expectation and precision for the residual error log-precision weights *λ*_*y*,*i*_. Specifically, we simulated BOLD responses for 100 synthetic subjects using a 5-region network models and both FFX and RFX modeling regimes (parameters per Supplementary Table 1). Then, we inverted chromatic DCMs under each hyperparameter setting, where the hyperprior mean and variance were varied as powers of 2, with integer exponents ranging from 2 to 7 and −7 to −3, respectively. Next, we conducted group-level inference over parameters 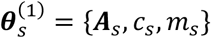, using BMA in the FFX case and PEB in the RFX case. We compared group-level MAP estimates to the ground truth, and the resulting RMSE surfaces over hyperparameter combinations were used to identify hyperpriors that optimized accuracy across both analysis approaches (Extended Data Fig. 1d–e).

### Face validity via in silico parameter recovery

To assess the identifiability of chromatic DCM parameters, we conducted simulations under both FFX and RFX assumptions. For each modeling regime, BOLD responses were generated in the time-domain (Eqs. 4– 5). In the FFX case, all synthetic subjects shared the same underlying model architecture and parameters. In the context of these analyses, the ground-truth effective connectivity matrix was generated as follows (using the same random seed for all *s* synthetic subjects):

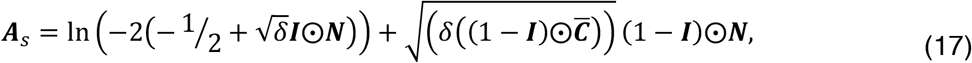

Here, 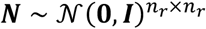 is IID noise, ⨀ represents element-wise multiplication, δ determines the variance of effective connectivity, and 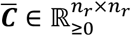 is a random symmetric (group-level structural connectivity) matrix. Note the log-normal sampling for intra-regional connections per their priors (Supplementary Table 2).

In contrast to the FFX case, in RFX simulations we permitted subject-specific deviations from a set of simulated group-level parameters. Namely, both effective connectivity and valency–spectra mapping parameters were treated as random effects, following:

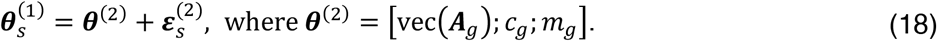

Here, 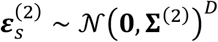 represents an independently seeded noise process for each synthetic subject, parametrized according to Eq. 14, *D* is the cardinality of ***θ***^(2)^, ***A***_*g*_ was obtained per Eq. 17, and *c*_*g*_ and *m*_*g*_ represent hand-coded group-level exponent mapping parameters. All other parameters (for example, the hemodynamic parameters ***θ***_*h*_) were fixed across simulations per FFX simulations (Supplementary Table 1).

To model realistic measurement conditions across both FFX and RFX simulations, power-law observation error was added to each simulated regional BOLD response. Specifically, for the *i*-th region, observation error *e*_*i*_(*t*) was generated by filtering white noise with a temporal kernel based on *g*_*e*,*i*,*i*_(*ω*), per the first line in Eq. 2. Each resulting region-specific error was normalized to unit variance and rescaled to match a desired SNR relative to the simulated BOLD signal *y*_*i*_(*t*), according to:

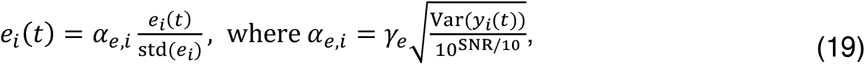

Although for consistency with Eqs. 1–2, Eq. 19 is written in terms of a continuous-time process, in the context of these simulations, the standard deviation and variance were computed over its discretized vector form.

### Hopf model-based validation of valency–spectra mapping

To assess whether a linear relationship between regions’ structural valency and the spectral slope emerges from a minimally coupled system, we simulated time series using a canonical network of coupled Hopf oscillators. Specifically, we employed a real-valued formulation of the Stuart–Landau oscillator (Hopf bifurcation normal form)^49^, with the system described by a 2*n*_*r*_-dimensional set of coupled differential equations:

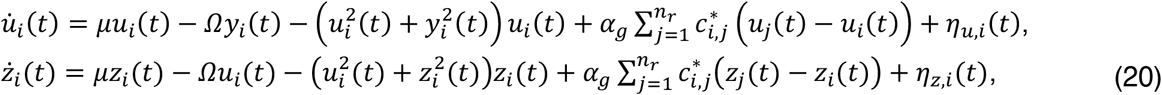

where *u*_*i*_(*t*) and *Z*_*i*_(*t*) denote the real and imaginary components of the oscillator states across the network, *μ* is the bifurcation parameter, *Ω* = 2*π*. 1/169 rad/s is the intrinsic angular frequency, 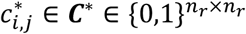 is a structural connectivity matrix (with no self-connections), and the parameter *α*_*g*_ ∈ ℝ_>0_ controls global coupling strength. Stochastic inputs *η*_*u*,*i*_(*t*) and *η*_*z*,*i*_(*t*), were drawn independently at each integration step from a zero-mean Gaussian 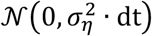, where *σ*_*η*_ denotes the standard deviation of dynamic noise. Here, we treated the real components of the oscillator states as the signal of interest.

Simulations were performed using the Euler–Maruyama integration method with a fixed integration step dt = 0.1, over *T* = 10^4^ time steps. Unless otherwise reported, simulations used *α*_*g*_ = 0.1, *μ* = −0.001, *σ*_η_ = 0.02, and connection density *d*_*c*_ = 1/10, with initial conditions drawn from a zero-mean Gaussian with standard deviation *σ*_0_ = 0.01. In addition to simulating dynamics with a large (*n*_*r*_ = 1000) random symmetric structural connectivity (Fig. 3b), we applied the model to an empirical connectome from the HCP dataset (*n*_*r*_ = 200, Fig. 3c).

To explore the role of criticality, we varied the bifurcation parameter *μ* ∈ [−1.5, 1.5] over 30 evenly spaced values. For each value of *μ*, we simulated the system 30 times with different noise realizations and computed two key metrics: the coefficient of determination (*R*^2^) for a linear model’s prediction of regional spectral exponents from structural valency, and metastability, defined as the standard deviation of the Kuramoto order parameter:

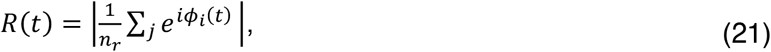

where ϕ_*i*_(*t*) is the instantaneous phase of *x*_*i*_(*t*)^50^. We reported 95% confidence intervals for both metrics, computed using the standard error of the mean across simulations. To estimate spectral exponents, PSDs were computed using Welch’s method, and linear regression was applied to the log-log spectrum over the 0.01 − 0.2 Hz frequency range with both log-frequency and log-power values were mean-centered.

### Construct validation against spectral dynamic causal modeling

To benchmark the chromatic DCM against an established model of resting-state effective connectivity, we compared its performance to spectral DCM as implemented in the SPM toolbox. As described in the Results section, both models were inverted using the same simulated datasets. All priors and inversion settings for spectral DCM followed the standard implementation.

### Data

We analyzed published and open-access neuroimaging and structural connectivity data for humans, macaques, marmosets, and mice. We note that analyses of macaque, marmoset, and mouse data were restricted to a single (average) hemisphere as determined by the tract-tracing and parcellation methodologies employed in the original studies. Essential methodological details concerning fMRI and structural connectivity are summarized below. Full procedures are described in the original sources.

### Human

Resting-state fMRI data were acquired for 200 healthy adults (ages 22–35 years, 100 female) from the HCP^20^ (for subject identifies per analysis, see Supplementary Table 4). All scans were acquired on a customized Siemens 3T Skyra scanner equipped with a 32-channel head coil and enhanced gradient system. Functional images were collected using a gradient-echo echo-planar imaging (EPI) sequence with multiband acceleration: TR = 0.72 s, echo time (TE) = 33.1 ms, flip angle = 52°, voxel size = 2 mm isotropic, and 1200 volumes. Here, in this study, analyses focused on the first, left-to-right phase-encoding direction run. Standardized preprocessing was applied using the HCP minimal preprocessing pipeline (v3.19.0)^51^, including independent component analysis-based denoising with FIX^52^, and time-series data were parcellated using a hybrid atlas comprising 200-regions from Schaefer and colleagues’ 17-network cortical atlas^53^, and Tian and colleagues’ 32-region subcortical atlas^54^. In analyses presented in Extended Data Fig. 1, regions for the hippocampal, striatal, and thalamic networks were specified based on those identified by Tian and colleagues as belonging to hippocampal, basal ganglia, and thalamic modules, respectively^54^.

Subject-level structural connectivity utilized in this study was derived from whole-brain probabilistic tractography applied to HCP subject’s diffusion-weighted MRI acquired using a spin-echo EPI sequence (TR = 5.52 s, TE = 89.5 ms, 1.25 mm isotropic voxels) with three b-value shells (1,000, 2,000, 3,000 s/mm^2^) and six b0 images (see Glasser and colleagues for all other acquisition and preprocessing details^51^). Following the MRtrix3 pipeline described by Arnatkeviciute and colleagues^55,56^, fiber orientation distributions were reconstructed in each participant’s native T1 space using the iFOD2 algorithm^57^, and Anatomically Constrained Tractography (ACT) was used to dynamic seed and generate 10 million streamlines per subject^58^. Streamlines were parcellated using the hybrid (Schaefer–Tian) atlas, and the resulting matrices were weighted by parcel surface area.

For network regions of interest **ℛ** ⊆ {1, . . ., *n*_*r*_}, normalized structural valency was computed as the column-wise sum of the symmetric, weighted structural connectivity matrix 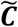, normalized by the maximum valency across the selected regions:

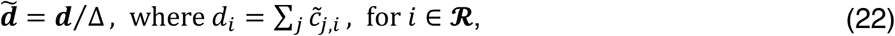

Note that this approach preserves the influence of whole-brain structural connectivity. Specifically, each region’s weight within the network reflects its total structural coupling with the rest of the brain (connections both outside and within the network). This is consistent with the idea that endogenous fluctuations emerge from a region’s embeddedness in the broader connectome.

### Macaque

Resting-state fMRI data were acquired for five adult male Macaca mulatta monkeys (ages 3.8–5.8 years) on a Siemens Tim Trio 3T MRI scanner and made available via the PRIMatE Data Exchange (PRIME-DE) project^34,35^. Animals were anesthetized using a dexmedetomidine–midazolam cocktail, with isoflurane gas used to maintain anesthesia during scanning. Functional images were acquired using a gradient-echo EPI sequence (TR = 2 s, TE = 29 ms, flip angle = 77°, voxel size = 1.5 × 1.5 × 2.5 mm^3^, 1200 volumes). Time series data were parcellated into 29 regions from Markov and colleagues’ 91-region atlas^59^, with the number regions determined by the number of injection sites in a related tract-tracing study^60^.

Data from this tract-tracing study were used to create the group-level (consensus) structural connectivity used here (Fig. 4f). Specifically, we utilized the 29-region, dense (edge-complete) connectivity matrix derived from the retrograde tracer data of Markov and colleagues^60^, made publicly available by Horvat and colleagues^29^. The asymmetric matrix entries encode the fraction of labeled neurons (FLN): the proportion of neurons in a source region projecting to a given target. Here, in the cross-species analyses (Fig. 4f), this matrix was symmetrized by averaging the upper and lower triangles, and the normalized structural valency was obtained per Eq. 22. In whole-brain analyses, we defined the structural valency in three distinct ways — summing either the rows (outgoing connections) of the asymmetric matrix, the columns (incoming connections) of the asymmetric matrix, or the columns of the symmetrized matrix — prior to normalization per Eq. 22.

### Marmoset

Resting-state fMRI data were acquired for 13 Macaca fascicularis monkeys (ages 2–4 years, 1 female) on a 9.4T Bruker horizontal scanner and made available via the Marmoset Brain Mapping (MBM) project^32^. Animals underwent a three-to four-week acclimatization protocol and were scanned while awake in the sphinx position with the head restrained using appropriate helmets. Functional images were acquired using a 2D gradient-echo EPI sequence (TR = 2 s, TE = 18 ms, flip angle = 70.4°, voxel size = 0.5 mm isotropic, 512 volumes). To align these data with structural connectivity, they were parcellated using the MBM projects’ implementation of the Paxinos’ atlas^61^.

Consensus structural connectivity for the marmoset analyses was derived from a publicly available, cellular-resolution tracer dataset comprising 143 retrograde injections registered to the Paxinos’ atlas by Majka and colleagues^30^. This yielded an asymmetric matrix encoding the FLN, which in the cross-species analyses (Fig. 4h), was symmetrized to compute the normalized structural valency per Eq. 22.

### Mouse

Resting-state fMRI data were acquired for 193 wild-type Mus musculus mice (ages 3–13 weeks, 82 female) on either a 9.4T or 11.75T small-animal MRI scanner equipped with a cryogenic surface phased-array coil, made available via the Radboud Data Repository^33,62^. All animals were anesthetized using a Mediso protocol, and ventilation was maintained throughout scanning. Functional images were acquired using a spin-echo and gradient-echo combined EPI sequence (TR = 1 s, TE = 9.2–15.0 ms, flip angle = 50° or 90°, voxel size = 200 *μ*m isotropic, 180 volumes). Standardized preprocessing steps were applied^33,62^, and time-series data were parcellated using a modified version of the Allen Mouse Brain Common Coordinate Framework (CCFv3) atlas, where dorsal and ventral anterior cingulate regions were aggregated to create the ACC parcel utilized in Fig. 4d.

The consensus structural connectivity utilized in this study was constructed by parcellating, according to our modified CCFv3 atlas, the voxel-resolution tract-tracing connectome made publicly available by Coletta and colleagues^31^. This 15,314-region connectome was constructed using a Voronoi-based resampling of a previously published voxel-wise-resolution connectome^63^, based on data from 428 anterograde viral tracer experiments conducted by Oh and colleagues available via the Allen Mouse Brain Connectivity Atlas (ABA)^64^. Per other species, this asymmetric structural connectivity was symmetrized for the cross-species analyses (Fig. 4i), and the normalized structural valency was computed per Eq. 22.

## Additional Information

### Data availability

Raw data analyzed in this study were sourced from published and open-access datasets. Derived data, including dynamic causal models, will be made available in summary form via a GitHub repository upon publication.

### Code availability

All simulations and model inversions were implemented using custom MATLAB scripts developed for this study. These scripts build on core routines from the Statistical Parametric Mapping (SPM12) toolbox. Except for data preprocessing executed prior to data acquisition for this study, all analyses were executed within the MATLAB environment. Code for the chromatic dynamic causal model (DCM), along with relevant analysis and visualization routines, will be made publicly via a GitHub repository upon publication.

## Acknowledgements

Data were provided by the Human Connectome Project, WU-Minn Consortium (Principal Investigators: David Van Essen and Kamil Ugurbil; 1U54MH091657) funded by the 16 NIH Institutes and Centers that support the NIH Blueprint for Neuroscience Research; and by the McDonnell Center for Systems Neuroscience at Washington University. M.D.G. is supported by an Australian Government Research Training Program Scholarship. M.D.G., L.N. and A.R. are funded by the Australian Research Council (ref. DP200100757). A.R. is also funded by Australian National Health and Medical Research Council Investigator Grant (ref. 1194910).

A.R. is affiliated with The Wellcome Centre for Human Neuroimaging supported by core funding from Wellcome (203147/Z/16/Z). A.R. is a CIFAR Azrieli Global Scholar in the Brain, Mind & Consciousness Programme.

## Author contributions

M.D.G. conceptualized the study. M.D.G., L.N., J.C.P., and A.R. designed the methodology. M.D.G. and J.C.P. performed the investigation and administered the project. M.D.G. developed visualizations. A.R. acquired funding. A.R., L.N. and A.F. supervised the project. M.D.G. wrote the original draft. All authors reviewed and edited the final manuscript.

## Extended Data

**Extended Data Fig. 1.**
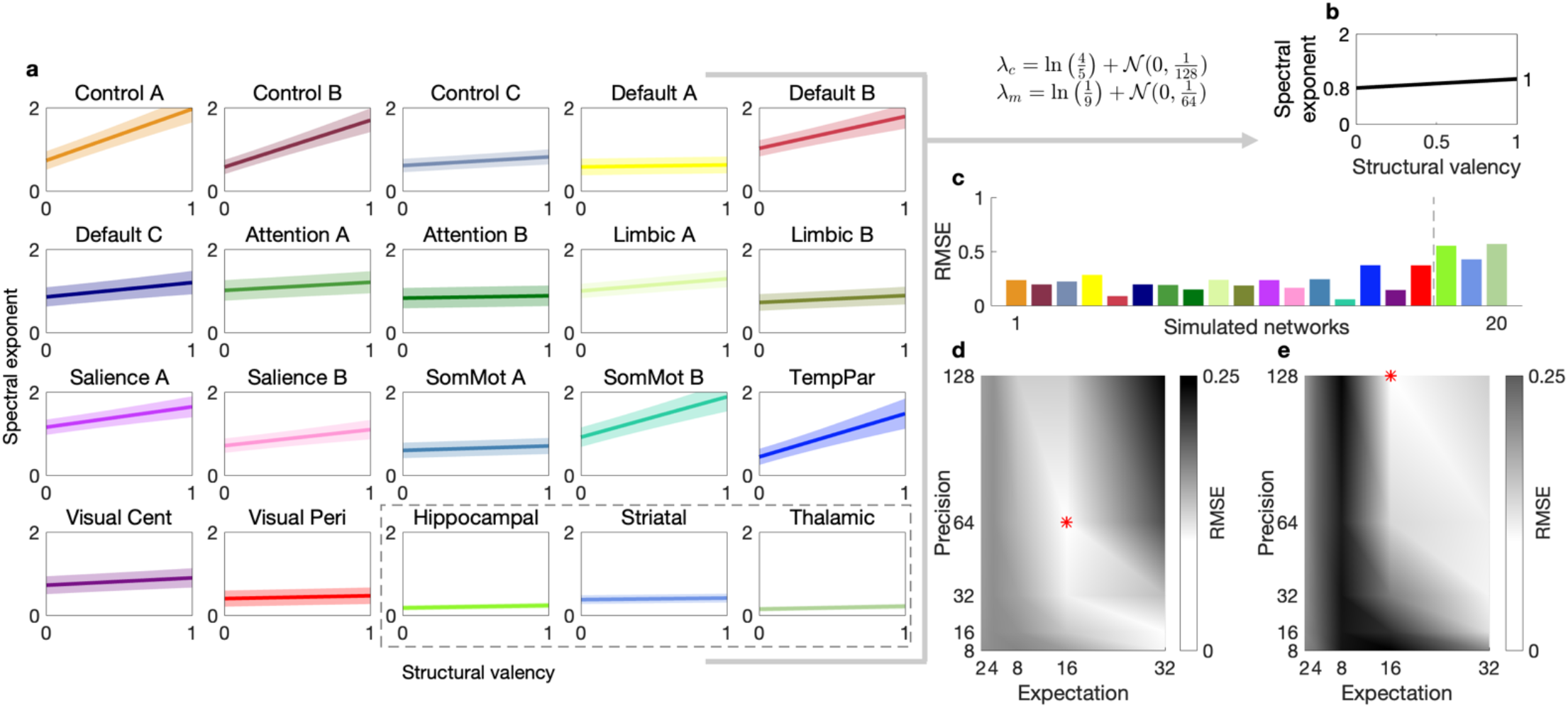
Establishing data-informed priors and hyperparameters in chromatic DCM. (**a**) Across 20 HCP-derived effective connectivity networks, group-level evidence-weighted PEB estimates revealed network-specific valency–spectra mappings, with stronger valency–exponent relationships in transmodal compared to unimodal and (in dashed box) subcortical networks. **(b)** The average mapping across networks provided empirical priors for *λ*_*c*_ and *λ*_*m*_ (controlling the slope and intercept of the valency–spectra mapping), used in all subsequent analyses. **(c)** RMSE between true and recovered parameters across 20 simulated networks confirmed good recoverability of valency–spectra mappings under the selected priors. **(d–e)** Grid search over prior expectation and precision values for residual error log-precision weights (*λ*_*y*,*i*_) revealed settings (red asterisks) that minimized RMSE across FFX **(d)** and RFX **(e)** simulations.

## Notes

### Competing Interest Statement

The authors have declared no competing interest.

## References

1. Huang, W. & Shu, N. AI-powered integration of multimodal imaging in precision medicine for neuropsychiatric disorders. Cell Reports Medicine 6, 102132 (2025).

2. Ji, J. et al. A Survey on Brain Effective Connectivity Network Learning. IEEE Trans. Neural Netw. Learning Syst. 34, 1879–1899 (2023).

3. The MICrONS Consortium et al. Functional connectomics spanning multiple areas of mouse visual cortex. Nature 640, 435–447 (2025).

4. Ereira, S., Waters, S., Razi, A. & Marshall, C. R. Early detection of dementia with default-mode network effective connectivity. Nat. Mental Health 2, 787–800 (2024).

5. Greaves, M. D., Novelli, L., Mansour L. S.,, Zalesky, A. & Razi, A. Structurally informed models of directed brain connectivity. Nat. Rev. Neurosci. 26, 23–41 (2025).

6. Razi, A. & Friston, K. J. The Connected Brain: Causality, models, and intrinsic dynamics. IEEE Signal Process. Mag. 33, 14–35 (2016).

7. Friston, K. J. Functional and Effective Connectivity: A Review. Brain Connectivity 1, 13–36 (2011).

8. Friston, K. J., Kahan, J., Biswal, B. & Razi, A. A DCM for resting state fMRI. NeuroImage 94, 396–407 (2014).

9. Razi, A., Kahan, J., Rees, G. & Friston, K. J. Construct validation of a DCM for resting state fMRI. NeuroImage 106, 1–14 (2015).

10. Bernal-Casas, D., Lee, H. J., Weitz, A. J. & Lee, J. H. Studying Brain Circuit Function with Dynamic Causal Modeling for Optogenetic fMRI. Neuron 93, 522-532.e5 (2017).

11. Almgren, H. et al. Variability and reliability of effective connectivity within the core default mode network: A multi-site longitudinal spectral DCM study. NeuroImage 183, 757–768 (2018).

12. Sultana, T. et al. Neural mechanisms of emotional health in traumatic brain injury patients undergoing rTMS treatment. Mol Psychiatry (2023) doi:10.1038/s41380-023-02159-z.

13. Murray, J. D. et al. A hierarchy of intrinsic timescales across primate cortex. Nat Neurosci 17, 1661– 1663 (2014).

14. Burt, J. B. et al. Hierarchy of transcriptomic specialization across human cortex captured by structural neuroimaging topography. Nat Neurosci 21, 1251–1259 (2018).

15. Fallon, J. et al. Timescales of spontaneous fMRI fluctuations relate to structural connectivity in the brain. Network Neuroscience 4, 788–806 (2020).

16. Sethi, S. S., Zerbi, V., Wenderoth, N., Fornito, A. & Fulcher, B. D. Structural connectome topology relates to regional BOLD signal dynamics in the mouse brain. Chaos: An Interdisciplinary Journal of Nonlinear Science 27, 047405 (2017).

17. Baria, A. T. et al. Linking human brain local activity fluctuations to structural and functional network architectures. NeuroImage 73, 144–155 (2013).

18. Gollo, L. L., Zalesky, A., Hutchison, R. M., Van Den Heuvel, M. & Breakspear, M. Dwelling quietly in the rich club: brain network determinants of slow cortical fluctuations. Phil. Trans. R. Soc. B 370, 20140165 (2015).

19. Moon, J.-Y. et al. Structure Shapes Dynamics and Directionality in Diverse Brain Networks: Mathematical Principles and Empirical Confirmation in Three Species. Sci Rep 7, 46606 (2017).

20. Van Essen, D. C. et al. The WU-Minn Human Connectome Project: An overview. NeuroImage 80, 62– 79 (2013).

21. Friston, K. J., Ashburner, J. T., Kiebel, S. J., Nichols, T. E. & Penny, W. D. Statistical Parametric Mapping: The Analysis of Functional Brain Images. (Elsevier Science, Burlington, 2011).

22. He, B. J., Zempel, J. M., Snyder, A. Z. & Raichle, M. E. The Temporal Structures and Functional Significance of Scale-free Brain Activity. Neuron 66, 353–369 (2010).

23. Fotiadis, P. et al. Structure–function coupling in macroscale human brain networks. Nat. Rev. Neurosci. 25, 688–704 (2024).

24. Mesulam, M. From sensation to cognition. Brain 121, 1013–1052 (1998).

25. Friston, K. J. et al. Bayesian model reduction and empirical Bayes for group (DCM) studies. NeuroImage 128, 413–431 (2016).

26. Penny, W. D. et al. Comparing Families of Dynamic Causal Models. PLoS Comput Biol 6, e1000709 (2010).

27. Deco, G., Kringelbach, M. L., Jirsa, V. K. & Ritter, P. The dynamics of resting fluctuations in the brain: metastability and its dynamical cortical core. Sci Rep 7, 3095 (2017).

28. Izhikevich, E. M. Dynamical Systems in Neuroscience: The Geometry of Excitability and Bursting. (The MIT Press, 2006). doi:10.7551/mitpress/2526.001.0001.

29. Horvát, S. et al. Spatial Embedding and Wiring Cost Constrain the Functional Layout of the Cortical Network of Rodents and Primates. PLoS Biol 14, e1002512 (2016).

30. Majka, P. et al. Open access resource for cellular-resolution analyses of corticocortical connectivity in the marmoset monkey. Nat Commun 11, 1133 (2020).

31. Coletta, L. et al. Network structure of the mouse brain connectome with voxel resolution. Sci. Adv. 6, eabb7187 (2020).

32. Tian, X. et al. An integrated resource for functional and structural connectivity of the marmoset brain. Nat Commun 13, 7416 (2022).

33. Grandjean, J. et al. Common functional networks in the mouse brain revealed by multi-centre restingstate fMRI analysis. NeuroImage 205, 116278 (2020).

34. Lv, Q. et al. Large-Scale Persistent Network Reconfiguration Induced by Ketamine in Anesthetized Monkeys: Relevance to Mood Disorders. Biological Psychiatry 79, 765–775 (2016).

35. Autio, J. A. et al. Towards HCP-Style macaque connectomes: 24-Channel 3T multi-array coil, MRI sequences and preprocessing. NeuroImage 215, 116800 (2020).

36. Tanner, J. et al. A multi-modal, asymmetric, weighted, and signed description of anatomical connectivity. Nat Commun 15, 5865 (2024).

37. Gilson, M., Moreno-Bote, R., Ponce-Alvarez, A., Ritter, P. & Deco, G. Estimation of Directed Effective Connectivity from fMRI Functional Connectivity Hints at Asymmetries of Cortical Connectome. PLoS Comput Biol 12, e1004762 (2016).

38. Greaves, M. D., Novelli, L. & Razi, A. Structurally informed resting-state effective connectivity recapitulates cortical hierarchy. Preprint at 10.1101/2024.04.03.587831 (2024).

39. Sokolov, A. A. et al. Asymmetric high-order anatomical brain connectivity sculpts effective connectivity. Network Neuroscience 4, 871–890 (2020).

40. Kim, J. H., Renden, R. & Von Gersdorff, H. Dysmyelination of Auditory Afferent Axons Increases the Jitter of Action Potential Timing during High-Frequency Firing. J. Neurosci. 33, 9402–9407 (2013).

41. Novelli, L., Friston, K. & Razi, A. Spectral Dynamic Causal Modelling: A Didactic Introduction and its Relationship with Functional Connectivity. (2023) doi:10.48550/ARXIV.2306.13429.

42. Stephan, K. E., Weiskopf, N., Drysdale, P. M., Robinson, P. A. & Friston, K. J. Comparing hemodynamic models with DCM. NeuroImage 38, 387–401 (2007).

43. Novelli, L., Friston, K. & Razi, A. Spectral Dynamic Causal Modelling: A Didactic Introduction and its Relationship with Functional Connectivity. Network Neuroscience 1–37 (2023) doi:10.1162/netn_a_00348.

44. Penny, W. D. & Roberts, S. J. Bayesian multivariate autoregressive models with structured priors. IEE Proc., Vis. Image Process. 149, 33 (2002).

45. Barrett, A. B. & Seth, A. K. Directed Spectral Methods. in Encyclopedia of Computational Neuroscience (eds. Jaeger, D. & Jung, R.) 1230–1234 (Springer New York, New York, NY, 2022). doi:10.1007/978-10716-1006-0_414.

46. Friston, K., Mattout, J., Trujillo-Barreto, N., Ashburner, J. & Penny, W. Variational free energy and the Laplace approximation. NeuroImage 34, 220–234 (2007).

47. Zeidman, P., Friston, K. & Parr, T. A primer on Variational Laplace (VL). NeuroImage 279, 120310 (2023).

48. Penny, W. D. et al. Comparing Families of Dynamic Causal Models. PLoS Comput Biol 6, e1000709 (2010).

49. Strogatz, S. H. Nonlinear Dynamics and Chaos. (CRC Press, 2018). doi:10.1201/9780429492563.

50. Breakspear, M., Heitmann, S. & Daffertshofer, A. Generative Models of Cortical Oscillations: Neurobiological Implications of the Kuramoto Model. Front. Hum. Neurosci. 4, (2010).

51. Glasser, M. F. et al. The minimal preprocessing pipelines for the Human Connectome Project. NeuroImage 80, 105–124 (2013).

52. Smith, S. M. et al. Resting-state fMRI in the Human Connectome Project. NeuroImage 80, 144–168 (2013).

53. Schaefer, A. et al. Local-Global Parcellation of the Human Cerebral Cortex from Intrinsic Functional Connectivity MRI. Cereb Cortex 28, 3095–3114 (2018).

54. Tian, Y., Margulies, D. S., Breakspear, M. & Zalesky, A. Topographic organization of the human subcortex unveiled with functional connectivity gradients. Nat Neurosci 23, 1421–1432 (2020).

55. Tournier, J.-D. et al. MRtrix3: A fast, flexible and open software framework for medical image processing and visualisation. NeuroImage 202, 116137 (2019).

56. Arnatkeviciute, A. et al. Genetic influences on hub connectivity of the human connectome. Nat Commun 12, 4237 (2021).

57. Tournier, J.-D., Calamante, F. & Connelly, A. Improved probabilistic streamlines tractography by 2nd order integration over fibre orientation distributions. Proc. Intl. Soc. Mag. Reson. Med. (ISMRM) 18, (2010).

58. Smith, R. E., Tournier, J.-D., Calamante, F. & Connelly, A. Anatomically-constrained tractography: Improved diffusion MRI streamlines tractography through effective use of anatomical information. NeuroImage 62, 1924–1938 (2012).

59. Van Essen, D. C., Glasser, M. F., Dierker, D. L. & Harwell, J. Cortical Parcellations of the Macaque Monkey Analyzed on Surface-Based Atlases. Cerebral Cortex 22, 2227–2240 (2012).

60. Markov, N. T. et al. A Weighted and Directed Interareal Connectivity Matrix for Macaque Cerebral Cortex. Cerebral Cortex 24, 17–36 (2014).

61. Paxinos, G. The marmoset brain in stereotaxic coordinates. Elsevier Academic Press (2012).

62. Grandjean, J. et al. Complex interplay between brain function and structure during cerebral amyloidosis in APP transgenic mouse strains revealed by multi-parametric MRI comparison. NeuroImage 134, 1–11 (2016).

63. Knox, J. E. et al. High-resolution data-driven model of the mouse connectome. Network Neuroscience 217–236 (2019).

64. Oh, S. W. et al. A mesoscale connectome of the mouse brain. Nature 508, 207–214 (2014).

