## Supplementary information for "Spectral imprint of structural embedding in effective connectivity"

**Table 1. | Parameters used in simulation-based analyses**

| Parameter | Description | Parameter value |
|---|---|---|
| $n_r$ | Number of regions | $\{5,7,10,12,15\}$ |
| $T$ | Number of time points | $1000$ |
| $TR$ | Repetition time | $1$ |
| SNR | Signal-to-noise ratio (dB) | $0$ |
| $d_c$ | Structural connectivity density | $1$ |
| $\delta$ | Variance of effective connectivity | $1/64$ |
| $c$ | Intercept of spectral exponent mapping | $4/3$ |
| $m$ | Slope of spectral exponent mapping | $1/4$ |
| $\alpha_v$ | Global amplitude of endogenous fluctuations | $10^{-5}$ |
| $\beta_e$ | Global spectral exponent of observation noise | $1/3$ |
| $\gamma_e$ | Normalization factor for observation error | $1$ |
| $k_h$ | Decay rate of the vasodilatory signal | $1$ |
| $\epsilon_h$ | Ratio of intra- to extra-vascular signal contributions | $1$ |
| $\tau_i$ | Transit times | $\sim \text{LogNormal}(0, e^{-8})$ |
| $\gamma_q$ | Second-level precision scale parameter | $\sim \mathcal{N}(0,1/16)$ |

Note: To ensure reproducibility, all simulations were initialized with a fixed random seed of 0 using MATLAB's 'rng' command. The number of regions $n_r$ varied across simulations according to the relevant design ($n_r = \{5,7,10,12,15\}$). For further details consult the Results section. For each simulation of $S$ instantiations (simulated subjects), the hemodynamic transit times $\tau_i$ were sampled independently for each region ($i = 1, \dots, n_r$) from a log-normal distribution, in addition to $\gamma_q$, sampled once per $S$ instantiations from a zero-mean Gaussian. This precision scale parameter $\gamma_q$, was only relevant in random-effects (RFX) simulations, where individual parameters represent deviations from a group-level distribution.

**Table 2. | Prior mean and variance for free parameters**

| Parameter | Description | Prior mean | Prior variance |
|---|---|---|---|
| $\ln(-2 \cdot a_{s,i,i})$ | Intra-regional effective connectivity | $-1/2$ | $1/64$ |
| $a_{s,i,j}$ | Inter-regional effective connectivity | $1/128$ | $1/64$ |
| $\ln \alpha_v$ | Global amplitude of endogenous fluctuations | $0$ | $1/64$ |
| $\ln \alpha_{e,i}$ | Global spectral exponent of observation noise | $0$ | $1/64$ |
| $\ln \beta_e$ | Normalization factor for observation error | $0$ | $1/64$ |
| $\ln \gamma_e$ | Decay rate of the vasodilatory signal | $0$ | $1/256$ |
| $\ln k_h$ | Ratio of intra- to extra-vascular signal contributions | $0$ | $1/256$ |
| $\ln \tau_i$ | Transit times | $0$ | $1/256$ |
| $\lambda_c$ | Latent intercept parameter for spectral mapping | $\ln 4/5$ | $1/128$ |
| $\lambda_m$ | Latent slope parameter for spectral mapping | $\ln 1/9$ | $1/64$ |
| $\lambda_{y,i}$ | Log-weights on precision of residual error | $16$ | $1/128$ |
| $\gamma_q$ | Second-level precision scale parameter | $0$ | $1/16$ |

Note: All priors are specified on the first-level chromatic dynamic causal model unless otherwise noted. Log-weights on the precision of residual error ($\lambda_{y,i}$) are treated as fixed hyperparameters that influence the likelihood but are not inferred during model inversion; that is, they do not yield posteriors. The second-level precision scale parameter $\gamma_q$ is relevant only in the context of second-level parametric empirical Bayes (PEB) models, where it modulates group-level shrinkage. All other priors used in second-level analyses are inherited directly from the first-level model. Namely, the prior distribution over the group-level parameters $\boldsymbol{\theta}^{(2)}$ is assumed to match the priors over the first subject's parameters $\boldsymbol{\theta}_1^{(1)}$, consistent with default PEB implementations in the statistical parametric mapping (SPM) toolbox.

**Table 3. l Homologous brain regions analyzed in cross-species analyses**

| Species | Common label | Region abbreviation | Region description |
|---|---|---|---|
| Human | V1 | VisCent_Striate_1 | Striate cortex |
| | V5 | VisPeri_StriCal_1 | Striate calcarine |
| | M1 | SomMotA_7 | Somatomotor A |
| | ACC | DefaultA_PFCm_1 | Medial prefrontal cortex |
| Macaque | V1 | V1 | Visual area 1 |
| | V5 | MT | Middle temporal area |
| | M1 | F1 | Frontal area F1 |
| | ACC | 24c | Area 24c |
| Marmoset | V1 | V1 | Visual area 1 |
| | V5 | V5 | Visual area 5 |
| | M1 | A4ab | Area 4ab |
| | ACC | A24d | Area 24d |
| Mouse | V1 | VISp | Primary visual cortex |
| | V5 | VISal | Anterolateral visual cortex |
| | M1 | MOp | Primary motor cortex |
| | ACC | ACAd | Dorsal anterior cingulate area |
| | ACC | ACAv | Ventral anterior cingulate area |

Note: Human data was parcellated using Schaefer and colleagues' 17-network, 200-region atlas (full parcel abbreviations use the prefix '17Networks_LH_', omitted for the sake of concision[1]). For macaques, regions correspond to those in Markov and colleagues' atlas[2]. Marmoset regions follow the Paxinos atlas[3,4], whereas mouse regions follow the Allen mouse brain connectivity atlas[5]. In our analyses, dorsal and ventral anterior cingulate areas were aggregated for the mouse.

**Table 4. | HCP S1200 participant identifiers used in analyses.**

| Analysis | Subject IDs |
|---|---|
| data-informed priors | 100408, 101107, 102311, 103010, 104012, 106016, 106521, 107018, 107321, 108222, 109830, 111716, 112112, 115320, 115724, 120212, 120414, 121618, 122317, 122620, 123521, 125222, 126325, 127630, 127832, 128026, 129028, 130114, 130417, 130518, 131217, 133019, 133827, 133928, 134425, 135730, 136732, 136833, 137633, 138534, 138837, 139233, 141422, 143830, 149539, 151021, 151526, 153025, 154229, 154734, 158338, 159239, 159946, 160830, 163129, 165840, 167036, 167440, 168947, 169343, 169444, 169747, 171330, 172938, 173334, 173435, 173839, 175439, 175742, 176037, 177241, 180735, 180836, 181131, 181232, 183741, 186141, 189652, 191841, 192136, 192237, 192641, 194443, 194645, 196750, 198249, 198451, 201111, 203923, 204420, 206323, 206525, 206929, 209329, 210011, 212116, 221319, 245333, 246133, 248339. |
| Cross-species analysis | 263436, 268850, 285446, 289555, 303119, 333330, 346137, 361234, 361941, 365343, 371843, 376247, 386250, 406836, 432332, 433839, 453441, 456346, 463040, 467351, 469961, 492754, 516742, 517239, 518746, 519647, 522434, 529549, 541640, 545345, 547046, 555954, 561242, 561444, 565452, 573249, 573451, 579867, 580650, 586460, 590047, 598568, 599671, 609143, 613538, 616645, 622236, 627852, 633847, 644246, 645450, 654552, 660951, 663755, 679568, 685058, 687163, 695768, 698168, 700634, 706040, 709551, 713239, 731140, 749058, 761957, 765056, 769064, 771354, 773257, 792564, 800941, 815247, 825048, 826454, 828862, 832651, 833148, 841349, 844961, 845458, 849264, 849971, 852455, 869472, 871762, 877168, 880157, 886674, 894774, 896879, 923755, 927359, 958976, 965771, 969476, 970764, 983773, 989987, 992673. |

Note: This table lists the anonymized participant identifiers from the Human Connectome Project (HCP) S1200 release included in analyses reported in the main text[6]. All data were used in accordance with the WU-Minn HCP Data Use Terms. Inclusion was based on the availability of both resting-state fMRI and diffusion-weighted imaging data

### References


1. Schaefer, A. *et al.* Local-Global Parcellation of the Human Cerebral Cortex from Intrinsic Functional Connectivity MRI. *Cereb Cortex* **28**, 3095–3114 (2018).
2. Markov, N. T. *et al.* A Weighted and Directed Interareal Connectivity Matrix for Macaque Cerebral Cortex. *Cerebral Cortex* **24**, 17–36 (2014).
3. Majka, P. *et al.* Open access resource for cellular-resolution analyses of corticocortical connectivity in the marmoset monkey. *Nat Commun* **11**, 1133 (2020).
4. Paxinos, G. The marmoset brain in stereotaxic coordinates. *Elsevier Academic Press* (2012).
5. Knox, J. E. *et al.* High-resolution data-driven model of the mouse connectome. *Network Neuroscience* **3**, 217–236 (2019).
6. Van Essen, D. C. *et al.* The WU-Minn Human Connectome Project: An overview. *NeuroImage* **80**, 62–79 (2013).
